# Overcoming tissue scattering in wide-field deep imaging by extended detection and computational reconstruction

**DOI:** 10.1101/611038

**Authors:** Yuanlong Zhang, Tiankuang Zhou, Xuemei Hu, Hao Xie, Lu Fang, Lingjie Kong, Qionghai Dai

## Abstract

Compared to the golden technique of point‐scanning multiphoton microscopy, line‐scanning temporal focusing microscopy (LTFM) is competitive in high imaging speed while maintaining tight axial confinement. However, considering its wide‐field detection mode, LTFM suffers from shallow penetration depth as a result of crosstalk induced by tissue scattering. In contrast to the spatial filtering based on confocal slit detection, we propose the extended detection LTFM (ED‐LTFM), the first technique to extract signals from scattered photons and thus effectively extend the imaging depth. By recording a succession of line‐shape excited signals in 2D and reconstructing signals under Hessian regularization, we can push the depth limitation in scattering tissue imaging. We valid the concept with numerical simulations, and demonstrate the performance of enhanced imaging depth in *in vivo* imaging of mouse brains.

---

Benefiting from its deep penetration, 3D sectioning capability, and low phototoxicity, etc, multiphoton microscopy (MPM) has found great applications in biomedical studies, including neuroscience and immunology [1, 2]. In conventional MPM, a tight focus is formed and thus the multi-dimensional imaging is generally performed by scanning the focus. However, the inertia of mechanical scanners limits the imaging speed[3, 4], which hampers the studies of most biological dynamics[5].

Recently, temporal focusing microscopy (TFM) has been proposed to achieve wide-field imaging while maintaining optical sectioning simultaneously [6, 7]: by introducing an angular dispersion to the excitation femtosecond pulses with a dispersion component, a spatiotemporal focus is formed when different frequency components overlap at the focal plane of the objective lens. Current progresses of TFM have demonstrated the confinement of two-photon wide-field excitation with decent axial resolutions[8]. Compared with the conventional point-scanning MPM, TFM enables high-speed imaging by parallel excitation [9, 10]. Generally, there are two modalities: planar-excitation TFM [11, 12] and line-scanning TFM [13]. In the former one, a planar region of the samples are excited in parallel; in the latter one, samples are excited by a sweeping line. In comparison, the axial focusing is weak in planar-excitation TFM, while LTFM exhibits better axial confinement and scattering resistance [14].

The well balance between imaging speed and axial resolution makes LTFM ideal for various applications, including laser processing [15] and large-scale imaging of biological dynamics [16]. To exploit the potential of LTFM in deep tissue imaging, Rowland *et al* has employing longer wavelength for minimizing the scattering suffered by the excitation beam[17]; we have proposed the focal modulation technique to modulate the excitation beam and thus eliminate the fluorescence background[18], and the hybrid spatio-spectral coherent adaptive compensation technique to compensate the aberrations experienced by the excitation beam[19].

Even though various strategies of multi-photon excitation reduce the effects of scattering for the excitation beam in TFM, as mentioned above, the crosstalk induced by scattering of the emitted fluorescence remains unsolved yet. Crosstalk of neighboring pixels in parallel readout via 2D sensors, such as sCMOS or EMCCD cameras, limits the signal-to-noise ratio (SNR) and the imaging depth in LTFM. To this end, confocal slit detection has recently been proposed [20] that exploits the same principle as that in confocal microscopy, where a confocal slit is conjugated to the line-shaped excitation. In practice, “virtual” confocal slit can be realized by setting the readout of sCMOS camera in “rolling shutter” mode, which would filter out the crosstalk in the direction orthogonal to the line. Apparently, such a technique could resist the scattering induced crosstalk between excitation lines effectively, but it would fail to resist the crosstalk along the excitation lines. Moreover, in confocal slit detection, one would lose most part of fluorescent signals when the scattering is severe.

Here we propose the ED-LTFM that could maintain the signal contrast and the resistance to scattering-induced noise in deep tissue imaging. A 2D fluorescent image is captured in each line-shape excitation position, so that the signals, including the scattering signals, are fully recorded. Then computational reconstruction is performed to recover the signals. Moreover, we incorporate Hessian regularization in the deconvolution, for the first time, which would ensure smooth transitions in the reconstructed images and thus reduce the artifacts caused by low SNR [21]. We demonstrate the enhanced performance of ED-LTFM in *in vivo* deep imaging of neurons in Thy1-YFP mouse brains and dynamic imaging of microglia in CX3CR1-GFP mouse brains.

As shown in Fig. 1(a), we denote one slice of the 3D sample as *f*(*x*, *y*), where (*x*, *y*) is the lateral coordinates. *f*(*x*, *y*) is excited column by column (along *x*-axis) by steering the laser line formed by temporal focusing. In conventional LTFM, the shutter of camera keeps open when the line-shape excitation beam excites the sample from one end to the other. At each excitation position, temporal focused laser line excites the sample and the emitted fluorescent signals go through the sample and the optical elements before being recorded by the camera. Unfortunately, the emitted fluorescent signals would suffer from tissue scattering, and photons from different excitation positions would mix in the sensor plane, as shown in Fig. 1(b). Consequently, the captured signals in wide-field detection (WD) LTFM thus could be written as

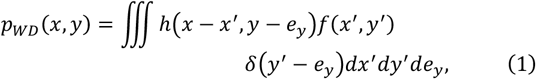

where *h* is the point spread function (PSF) of the system, *e*_y_ is the location of the line-shape excitation beam. The captured image *p*_WD_(*x*, *y*) would suffer from serious crosstalks along both *x* and *y* axis if *h* is largely affected by tissue scattering.

**Fig. 1.**
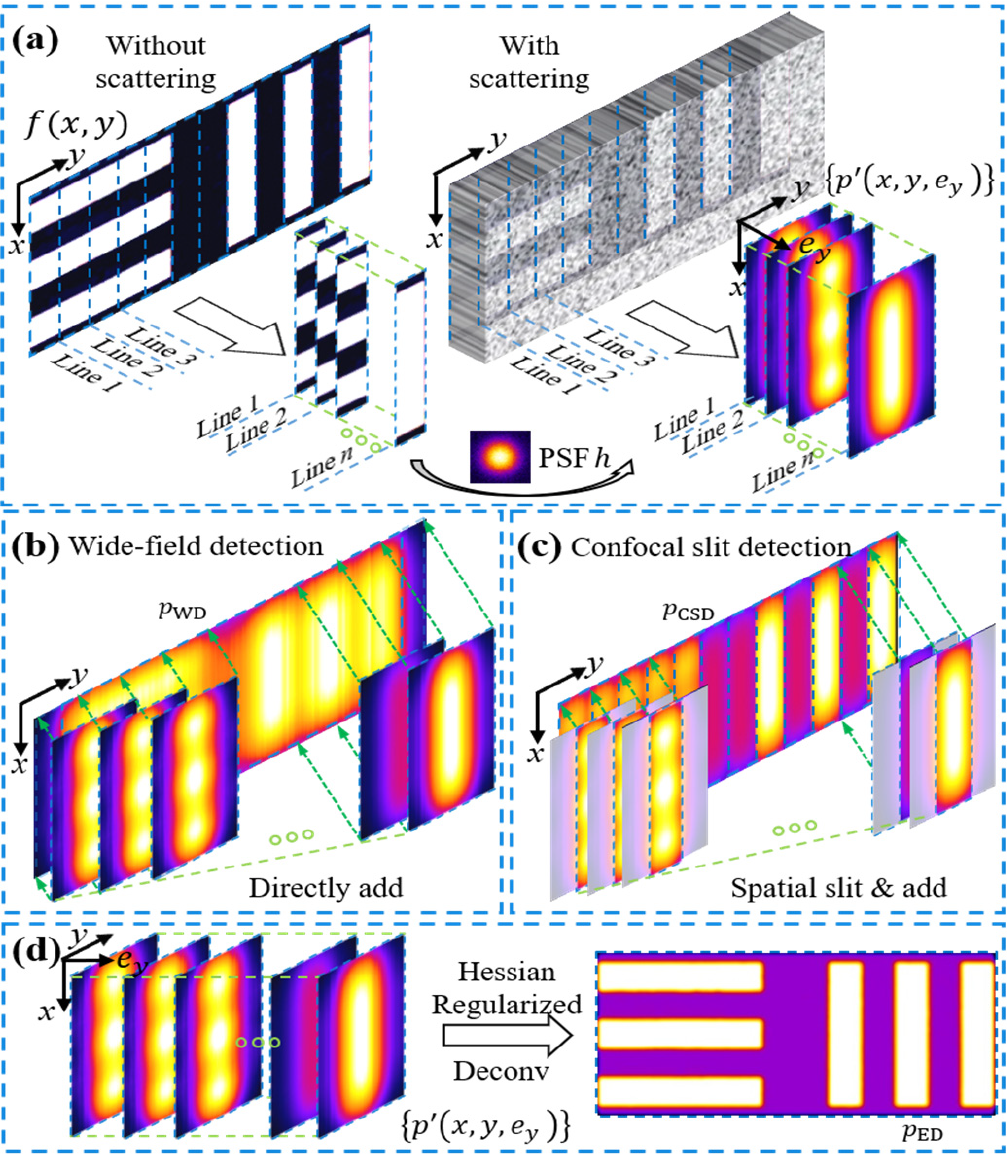
Illustrations of the ED-LTFM method. (a) The scattering PSF *h* makes the excitation line-shape signals overlapped along *y*-axis (direction of scanning) and blurred along x-axis (direction of excitation line). (b) Line-shape signals in conventional LTFM are integrated by the wide-field detection camera, thus are overlapped in both *x*- and *y*-axis. (c) A confocal slit is inserted before the sensor, which can reduce the crosstalk along *y*-axis but not the crosstalk along *x*-axis. (d) The extended detection (ED) technique records all of the signals, including scattering signals, for subsequent computational reconstruction, which could reduce cross talk along both along *x*- and *y*-axis.

For confocal slit detection (CSD), a detection slit is inserted before the camera to block the scattering photons. The captured signals by CSD then could be written as

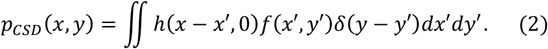

Even though the crosstalk along *y*-axis could be effectively reduced by the confocal slit detection, the crosstalk along *x*-axis remains, as shown in Fig. 1(c). Moreover, if scattering enlarges the PSF ℎ, the confocal slit would cut a majority of the signals and affect the final imaging SNR.

To suppress the effect of scattering along both *x*-axis and *y*-axis, we propose the ED-LTFM method that fully utilizes the crosstalk information to recover the original signals. More specifically, ED-LTFM acquires an image stack instead of a single image, for a single layer imaging:

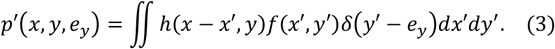

In other words, {*p*′(*x*, *y*, *e*_*y*_)} is the set of images that are excited by the laser line at each excitation position, as shown in Fig. 1(d). To recover image *f*(*x*, *y*) from the captured {*p*′(*x*, *y*, *e*_*y*_)}, we propose the following optimization problem:

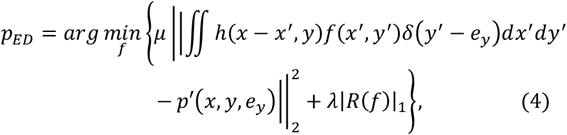

where

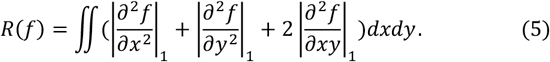

The penalty function *R*(*f*) used here is the Hessian regularization, *μ* and λ are the weights of losses. We adopt the alternative direction method of multipliers (ADMM) [22] to solve Eq. (4). Note that it is necessary to firstly estimate the PSF *h*. We estimated the PSF by fitting the scattering pattern of infinitesimal source with Henyey-Greenstein (H-G) function [16]. To summarize, the reconstruction works by iteratively updating two steps: (1) PSF estimation by fitting isolated scattering pattern in {*p*′(*x*, *y*, *e*_*y*_)} with H-G function, and (2) Hessian regularized deconvolution with the obtained PSF.

After setting up the algorithm, we evaluate the performance of the proposed methods via numerical simulations (Fig. 2). We show the retrieved images of the microtubules by WD, CSD and ED, in Figs. 2(b), 2(c) and 2(d), respectively. We split the original microtube image into columns, then convolve each column with the PSF *h* to generate {*p*′(*x*, *y*, *e*_*y*_)}. A 40 dB Gaussian white noise is added to simulate the real experiments. After generating {*p*′(*x*, *y*, *e*_*y*_)}, *p*_WD_, *p*_CSD_ and *p*_ED_ are then calculated via Eqs. (1), (2), and (4), respectively. We could see that *p*_ED_ shows the lowest background among *p*_WD_, *p*_CSD_ and *p*_ED_. We quantitatively measure the width of retrieved microtubules achieved with these three methods and could see that *p*_ED_ resembles ground truth the most, which demonstrates great advantages of our ED-LTFM in strong scattering and noisy conditions. To compare the retrieval quality quantitatively, we labeled the structured structural similarity index (SSIM) of each method in Figs. 2(b), 2(c) and 2(d) [23]. The result with ED is 2.1 times better than that with CSD and 15.5 times better than that with WD in terms of the index value.

**Fig. 2.**
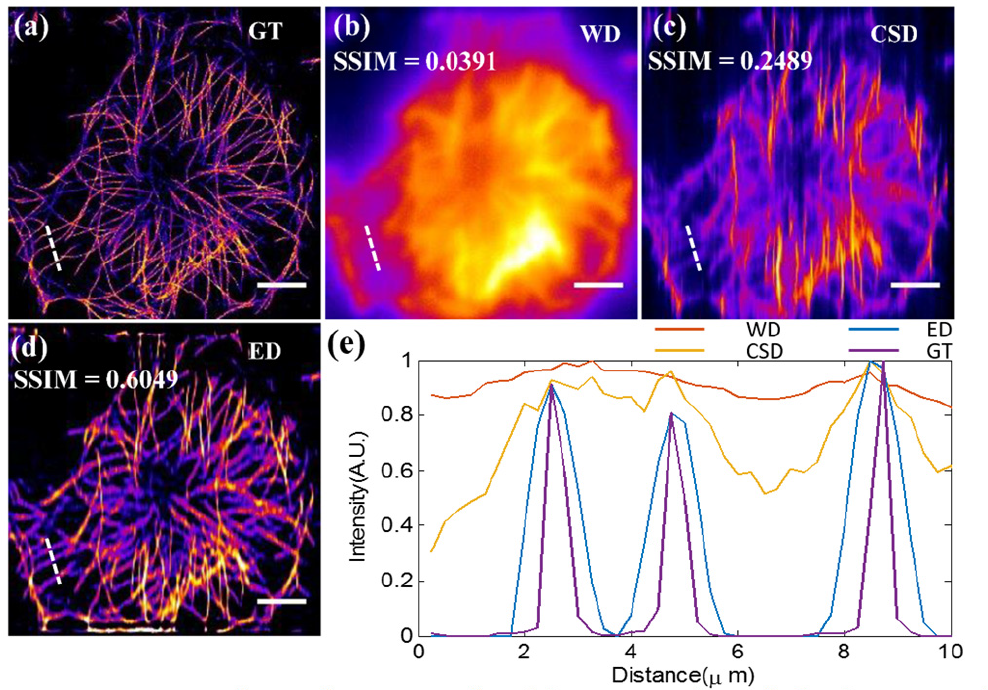
Numerical simulation results. (a) A ground-truth (GT) image of microtubule used in simulations. (b)(c)(d) Images retrieved with WD, CSD and ED, respectively. (e) The intensity along dashed lines in (a)(b)(c)(d). Upper left values: SSIM index compared to GT. Scale bar: 10 μm.

We then demonstrate the effectiveness of ED-LTFM through experiments. Fig. 3 shows the optical configuration of ED-LTFM. We use an 80 MHz, ∼120 fs laser (Chameleon Discovery, Coherent) for two-photon excitation at 920 nm, and a following electro-optical modulator (M3202RM, Conoptics) to control the laser intensity. The laser beam is expanded to ~ 5 mm with a telescope (L1: f =60 mm, L2: f =150 mm), and then scanned in the vertical direction with a one-dimensional galvanometer (GVS211, Thorlabs). The beam is focused to a thin line on the surface of the diffraction grating (Edmund Optics, 830 lines/mm) with a cylindrical lens (f = 300 mm). The incident angle to the grating is ∼50° to ensure that the central wavelength of the 1st diffraction light is perpendicular to the grating surface. The spectrally-spread pulse is collimated with a collimating lens (L3: f =200 mm), so that the expanded beam fulfills the back pupil of the objective (25×, 1.05 NA, water immersion, Olympus, XLPLN25XWMP2). A line-shaped laser beam, around 80 µm in length, is formed at the focal plane of the objective. An epi-fluorescence setup is built-up for image acquisition, including a dichroic mirror (DMSP750B, Thorlabs), a bandpass filter (E510/80, Chroma), a 200 mm tube lens (L4, TTL200-A, Thorlabs), and an sCMOS (Zyla 5.5 plus, Andor). Three-dimensional imaging is performed by axially moving the sample stage (M-VP-25XA-XYZL, Newport).

**Fig. 3.**
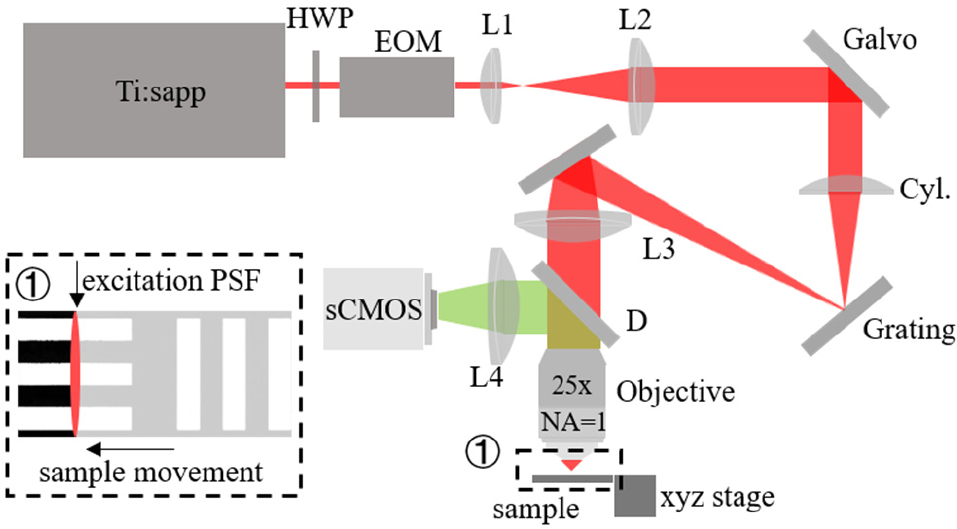
Optical configuration of the ED-LTFM. EOM, electro-optical modulator; HWP, half wave plate; Cyl. Lens, cylindrical lens; D, dichroic mirror. Inset 1: In our setup, the excitation line is fixed and the sample stage moves to achieve line scan.

In wide-field detection *p*_WD_, the camera keeps open during the line-shaped beam scans the sample. To capture {*p*′(*x*, *y*, *e*_*y*_)}, we park the beam in the center of the field-of-view (FOV) and translate the sample stage to finish the 1D scan, which will simplify the experimental setup for confocal slit detection and make the imaging field unlimited by the FOV of the objective. In this case, the camera captures a 650-by-200-pixel image at each stage position and the total {*p*′(*x*, *y*, *e*_*y*_)} stack is formed by 400 captures. To mimic the confocal slit detection, we use the center signal of the captured image stack {*p*′(*x*, *y*, *e*_*y*_)} to recover *p*_CSD_(*x*, *y*). *p*_ED_ is calculated from the proposed reconstruction algorithm. For fair comparison, the exposure time of each row in all three cases are the same (50ms). Note that the excitation line is along *x* axis, the same as the derivation above.

We first demonstrate the performance of ED-LTFM in *in vivo* imaging of living Thy1-YFP mice (JAX No. 005582). After craniotomy, we conduct acute imaging of neurons in the cerebral cortex with the living mice under anesthesia[24] (All procedures involving mice were approved by the Animal Care and Use Committees of Tsinghua University). In Figs. 4(a), 4(b) and 4(c), we show the maximum intensity projection (MIP) of neurons along the *z*-axis of a 13-µm-thick image stack (80–92 μm under the dura) acquired via WD, CSD and ED, respectively. For precise comparison, we show the zoomed-in view of the captured images in Figs. 4(d), 4(e), and 4(f). As expected, the dendrites are blur in WD due to the strong scattering, while CSD techniques help to eliminate the crosstalk induced by scattering along *y*-axis, and ED effectively eliminates the crosstalk along both *x*-axis and *y*-axis. In Fig. 4(j), we show the signal improvement of ED over that in WD and CSD by quantitatively comparing intensity along the dashed line in Figs. 4(d), 4(e) and 4(f). In Fig. 4(k), we also show that the proposed ED technique could help improve the signal contrast along *z*-axis via MIP along the *y*-axis of a 10 μm-thick *x‐z* image stack [labeled by the dashed box in Figs. 4(a), 4(b) and 4(c)] in Figs. 4(g), 4(h) and 4(i). It can be seen that the improvements of ED-LTFM are obvious.

**Fig. 4.**
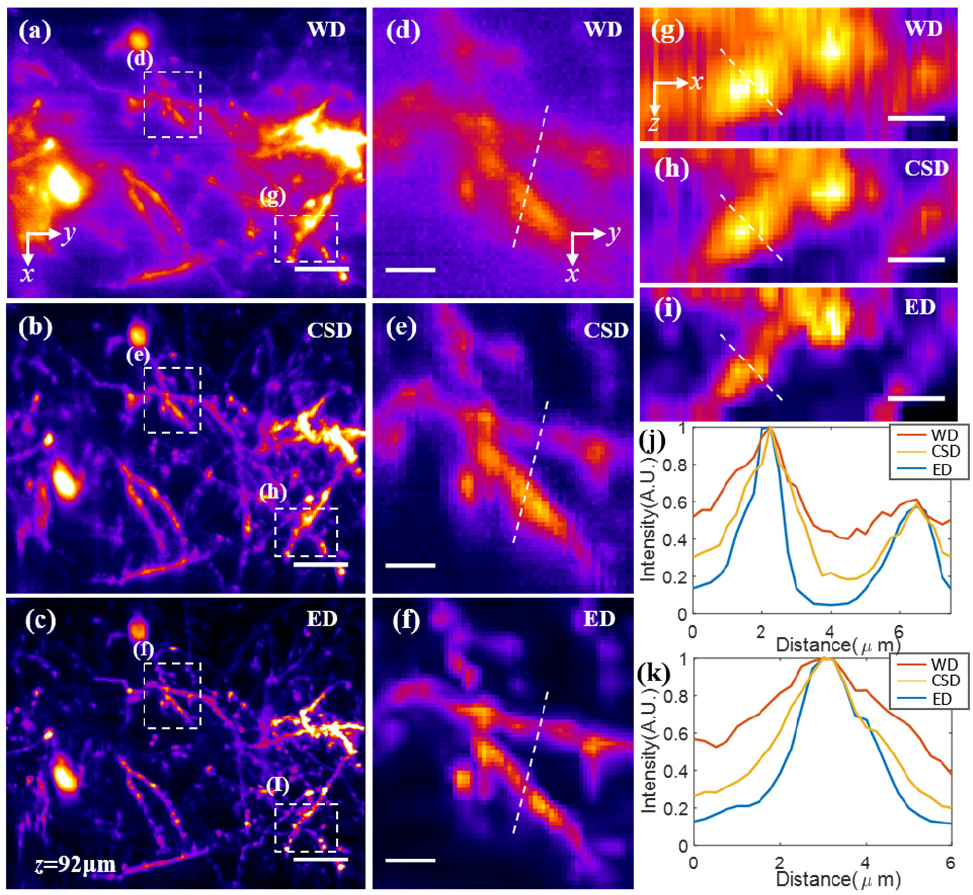
Comparison of different techniques in *in vivo* deep imaging of neurons. (a)(b)(c)MIPs of neurons along z-axis of a 13-μm-thick image stack (80–92 μm under the dura) acquired with the WD, CSD and ED, respectively. (d)(e)(f) Zoom-in views of lateral area marked by the dashed box in (a), (b) and (c), respectively. (g)(h)(i) MIPs along y-axis of a 10 μm thick x-z stack [marked by the dashed box in (a)(b)(c)]. The MIPs of x-z stacks are shown with bilinear interpolation along z-axis to equate the lateral and axial pixel sizes. (j) Intensity profiles along the indicated lines in (d)(e)(f). (k) Intensity profiles along the indicated lines in (g)(h)(i). Scale bars in (a)(b)(c) are 10 μm, in (d)(e)(f) are 3 μm.

Furthermore, we demonstrate the performance of ED-LTFM in *in vivo* dynamical imaging of microglia cells in living CX3CR1-GFP mice (JAXNo. 003782). In Figs. 5(a), 5(b) and 5(c), we show the MIP along the *z*-axis of a 7-µm-thick image stack acquired via WD, CSD, and ED, respectively, at the depths of 172–178 μm under the dura. To the best of our knowledge, no such penetration depths in LTFM has been demonstrated *in vivo* before. We could see that the background noise by ED has been suppressed significantly compared to that by both WD and CSD. By selecting a small part of Figs. 5(a), 5(b) and 5(c), we could see that CSD eliminates the crosstalk along *y*-axis effectively but fails to eliminate the cross talk along *x*-axis. The fine process could be recovered by ED effectively, which even could not be resolved in original WD images. We also image the “non-resting” dynamical movement of microglia cells over a total time of 16 minutes within a depth range of 30 μm. In Figs. 5(g), 5(h) and 5(i), we can see that, compared with the results from WD and CSD, ED can record the movement of the processes of microglia cells with fine details.

**Fig. 5.**
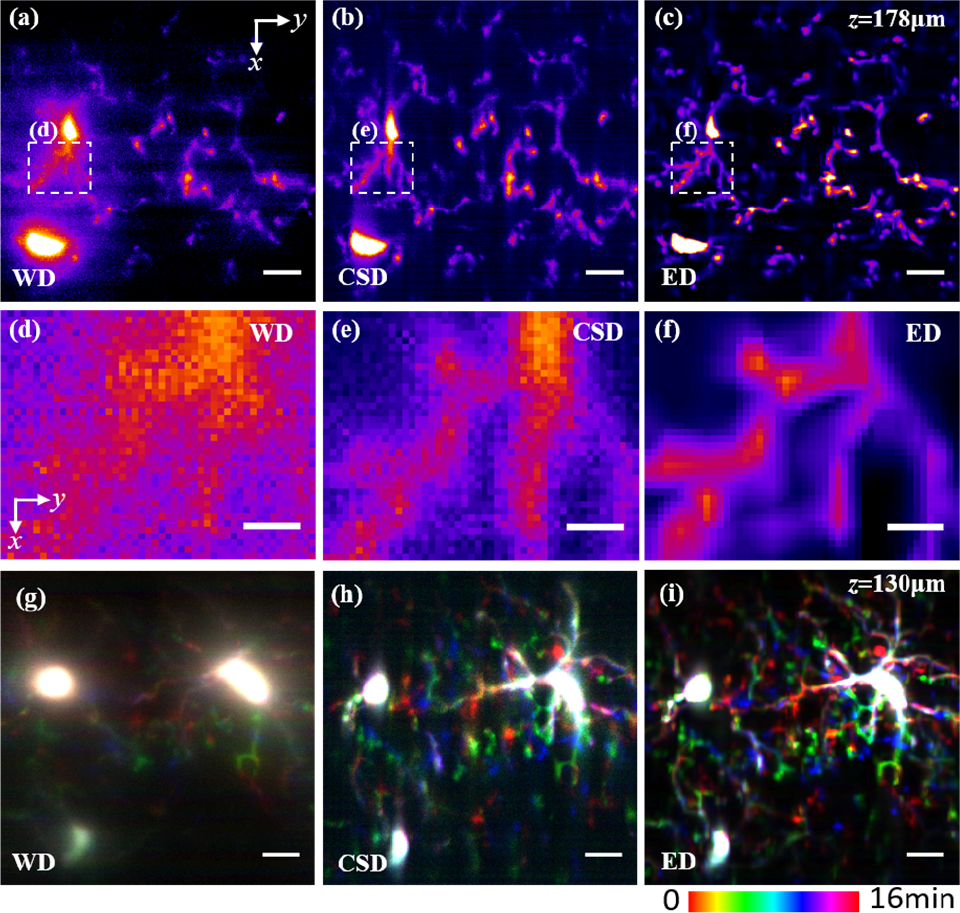
Comparison of different techniques in *in vivo* deep imaging of microglia cells and their dynamics. (a)(b)(c) MIP of microglia cells along *z*-axis of a 7-μm-thick image stack (172–178 μm under the dura) acquired with the WD, CSD and ED, respectively. (d)(e)(f) Zoom-in views of lateral area marked by the dashed box in (a), (b) and (c), respectively. (g)(h)(i), temporal color-coded MIP sequences of microglia cells along 30-μm-thick image stack (100–130 μm under the dura) acquired with WD, ED and CSD. Scale bars in (a)(b)(f)(g)(h)(i) are 10 μm, in (d)(e)(f) are 3 μm.

In summary, we propose a novel technique for overcoming tissue scattering in wide-field deep imaging by extended detection and computational reconstruction. Through both numerical simulations and *in vivo* imaging experiments, we have demonstrated that the proposed ED-LTFM can effectively enhance the penetration depth. For further improvement, a femtosecond laser of low repetition rate but high pulse energy would benefit for larger penetration depth and higher SNR. It is worth noting that ED-LTFM can also be integrated with other strategies, such as three-photon LTFM[17] and adaptive optics based LTFM[19], to push the limits of imaging depth in scattering tissues.

## Acknowledgment

YZ thanks Yingjun Tang for helps in sample preparations. LK thanks the support from Tsinghua University and the “Thousand Talents Plan” Youth Program.

